# Construction of an MRCUT1 Cutinase-Expressing *Saccharomyces boulardii* Probiotic Yeast Strain Capable of Degrading Polyethylene Terephthalate (PET) Microplastics

**DOI:** 10.1101/2025.07.16.665244

**Authors:** Luisa Marie S. Jingco, Albert Derrick R. Recio, Addie Franchesca D. Manrique, Daniel Jericho H. Dungo, Paul Cedric Bernardo, Gabriel Mendoza, Raphael Willard Mosquito, Lexi Caitlin Murillo, Joe Anthony H. Manzano, O.P. Nicanor Austriaco

## Abstract

Excessive use of plastics has led to an unprecedented global accumulation of plastic waste, particularly of the polyethylene terephthalate (PET) polymer, resulting in severe environmental and health challenges due to the proliferation of microplastic particles. To respond to this environmental challenge, especially in the Philippines, where single-use plastic sachet pollution is rampant, we genetically engineered *Saccharomyces boulardii* yeast cells to express and secrete the cutinase enzyme (MRCUT1) derived from the fungus *Moniliophthora roreri*. Our results suggest that our engineered yeast can degrade pre-weighed PET fragments over a twenty-one-day period *in vitro,* with significant measurable weight reductions observed within the first week of incubation. Statistical analysis (One-way ANOVA, *p* = 0.0112) confirmed that our MRCUT1 engineered strain degraded PET at a significantly higher rate than non-engineered control cells. To the best of our knowledge, this is the first time that a cutinase capable of PET degradation has been successfully expressed in and secreted by yeast. It also opens up the possibility that our MRCUT1-expressing probiotic could be used prophylactically to mitigate the microplastic bioaccumulation in the gastrointestinal tracts of common food fish like the rabbitfish (*Siganus fuscescens*) and the milkfish (*Chanos chanos*), both of which are popular aquaculture fish in the Philippines.

## INTRODUCTION

Plastics have become indispensable due to their low cost, durability, and versatility [1,2], leading to widespread use in packaging, transportation, and textiles. The demand for plastic resulted in over 359 million tons of plastic products produced in 2018, with polymers like polyethylene terephthalate (PET), polyvinyl chloride (PVC), polypropylene (PP), and polystyrene (PS) being among the most prevalent [2,3]. PET alone accounted for 30.5 million tonnes in 2019 and continues to grow in usage [4]. Of the total plastic waste, around 9% is recycled, 12% is incinerated, and 79% is disposed of in the environment [5].

Microplastics, which are plastic particles of 1μm to 5μm in size, have become ubiquitous in the environment. Several studies confirmed the presence of these particles in the atmosphere, aquatic habitats, and organisms. Bioaccumulation has been reported in the gastrointestinal tracts of invertebrates [6], turtles [7], and fishes [8]. These particles pose increasing risks to ecological and human health [9,10,11,12]. Microplastic exposure has been associated with various adverse effects, including insulin resistance, weight gain, and microbial contamination, underscoring the urgent need for effective mitigation strategies [13,14,15,16,17,18].

The Philippines is a major contributor to global ocean plastic pollution, contributing around 360 metric tons of plastic waste to the oceans every year [19]. A significant portion of this waste comes from single-use plastic sachets, often made of polyethylene terephthalate (PET), which are commonly sold in the “sari-sari” convenience stores that dot the archipelago of 7,641 islands [20]. The World Bank estimates that Filipinos use 163 million plastic sachets daily [21], a staggering number that leads to widespread microplastic pollution in the marine environment. Studies have shown significant microplastic contamination in the gastrointestinal tracts of common food fish, like the rabbitfish (*Siganus fuscescens*) and the milkfish (*Chanos chanos*), both of which are popular in the Philippines [22,23]. The management of plastic pollution, especially single-use plastic sachet pollution, is now of significant political and social concern in the country.

Microbial degradation is considered a promising strategy for bioremediation to address the growing problem of plastic pollution [24]. For waste management of plastic sachets and commercial bottles made of polyethylene terephthalate (PET), both natural and genetically modified microorganisms that degrade the PET polymer are available [25]. Natural microbes include bacteria like *Ideonella sakaiensis,* which can degrade up to 75% of PET fragments at 28°C over 70 days [26], and fungi like *Aspergillus niger*, which can degrade 53% of PET fragments in a month [27]. Engineered microbes, primarily *Escherichia coli*, expressing a range of enzymes, including cutinases and PETases, could degrade up to 35% of PET in a month [25]. Surface display of a PETase on *E. coli* resulted in cells that could degrade PET films [28]. Overexpression of the PETase from *Ideonella sakaiensis* in the yeast, *Yarrowia lipolytica*, results in PET film degradation [29]. In contrast, it is striking that the expression in the bacteria, *Escherichia coli,* of a PET hydrolytic enzyme from *Caldimonas taiwanensis* termed *Ct*PL-DM led to active PET degradation, but not when it was expressed in the yeast, *Pichia pastoris* [12].

MRCUT1 is a promising variant of the cutinase protein family that is derived from the phytopathogenic fungus *Moniliophthora roreri*. Vázquez-Alcántara et al. (2021) [30] expressed the mrcut1 gene in *E. coli* and reported around 31% PET microplastic weight degradation after 21 days in a reconstituted enzyme solution. To provide an alternative strategy for bioremediation, we decided to engineer the probiotic yeast, *Saccharomyces boulardii*, to express and secrete the MRCUT1 for the external degradation of PET microplastics. To the best of our knowledge, this is the first time that a cutinase capable of PET degradation has been successfully expressed and secreted in yeast. It also opens up the possibility that our MRCUT1-expressing probiotic yeast could be used prophylactically to mitigate the microplastic bioaccumulation in the gastrointestinal tracts of common food fish, like the rabbitfish (*Siganus fuscescens*) and the milkfish (*Chanos chanos*), both of which are popular in the Philippines. Milkfish, known locally as “bangus,” is a cornerstone fish species of the aquaculture industry in the country.

## RESULTS AND DISCUSSION

### Construction of an MRCUT1 Cutinase-Expressing *Saccharomyces boulardii* Yeast Strain

The mrcut1 gene, a promising variant of the cutinase gene family that is derived from the phytopathogenic fungus *Moniliophthora roreri*, was previously expressed in *E. coli* and was associated with a 31% PET microplastic weight degradation loss for a 21-day period by a purified enzyme solution obtained from the recombinant bacteria [30]. The same study suggested that the MRCUT1 enzyme contains a catalytic triad composed of Ser120-Asp172-His185, essential for ester bond hydrolysis. Enzyme activity assays indicate that MRCUT1 functions optimally at temperatures between 30°C and 40°C and under mildly alkaline conditions. However, it is unclear from this study if the MRCUT1 enzyme was secreted into the environment since the degradation studies were done with enzyme solutions, not intact recombinant cells.

To provide an alternative bioremediation strategy that could be deployed in environments polluted with PET microplastics, we decided to create an MRCUT1-expressing *Saccharomyces boulardii* that could secrete the recombinant enzyme into its surroundings. Moreover, since *S. boulardii* is a probiotic yeast considered Generally Recognized as Safe [31], we speculate that our MRCUT1-expressing yeast could be used prophylactically to counteract microplastic accumulation in the gut of common food fish, like the rabbitfish (*Siganus fuscescens*) and the milkfish (*Chanos chanos*), both of which are popular aquaculture fish in the Philippines.

We transformed the *S. boulardii* M2 strain [32] with a high-copy plasmid expressing the mrcut1 gene tagged with 6xHis with the yeast mating factor secretion signal at its N-terminus. To confirm the expression of MRCUT1 in our transformed cells, we performed Western Blot analysis using an anti-6xHis monoclonal antibody. As shown in Figure 1, our results revealed distinct bands in extracts obtained from independently transformed yeast strains containing our plasmid that were absent in cell extracts of untransformed yeast. We infer that the 25-34 kDa upper band corresponds to the predicted molecular mass of MRCUT1 fused with the yeast α-mating factor signal peptide (30.1 kDa). A lighter band, observed in the 20-24 kDa range, represents the cleaved, mature form of MRCUT1 (21 kDa), which remains after the yeast mating factor secretion signal is cleaved by the Kex2p endopeptidase [33].

**Figure 1.**
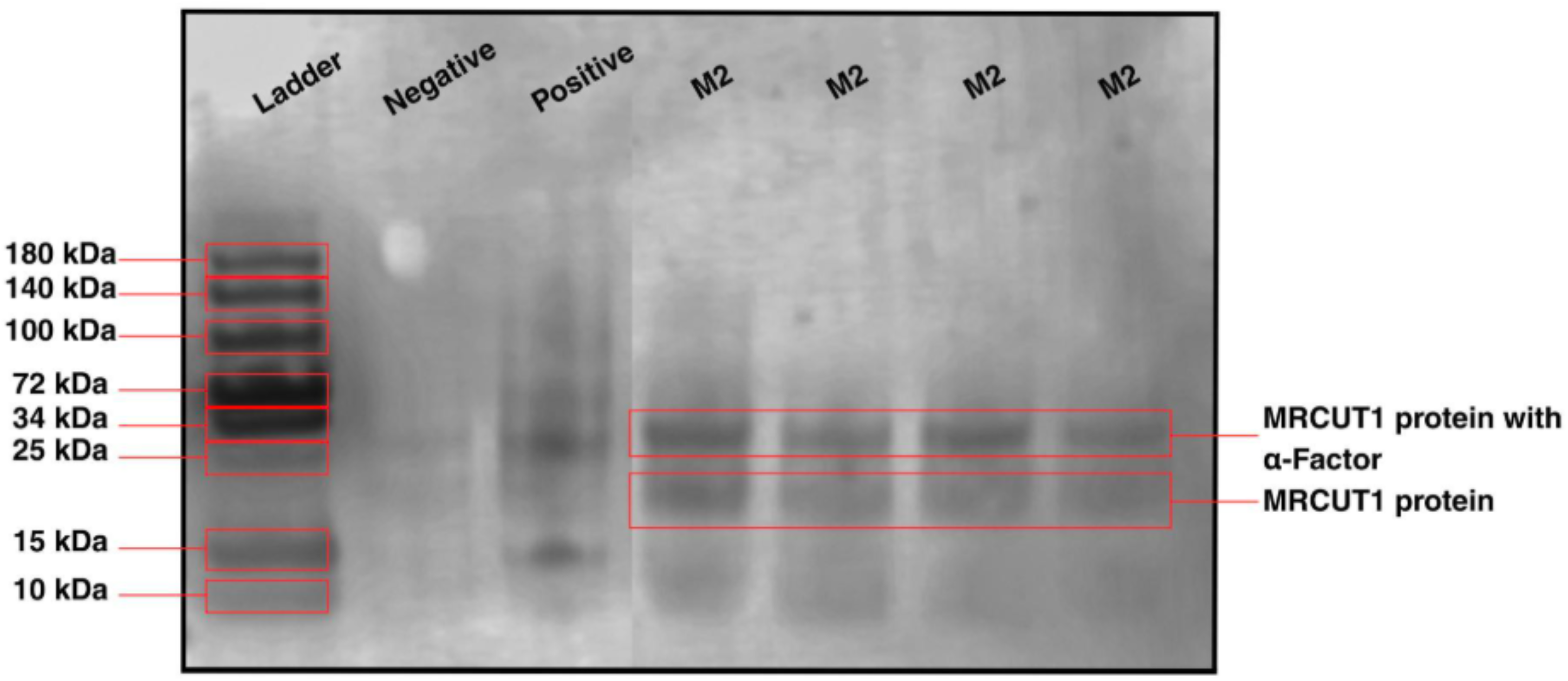
Expression of MRCUT-1 in Recombinant *Saccharomyces boulardii*. Lane 1: Protein Ladder (10–180 kDa). Lane 2: Native M2 strain (negative control); Lane 3: Recombinant M2 strain secreting SARS-CoV-2 RBD (positive control); Lanes 4–7: Recombinant M2 strains secreting MRCUT1.

### Our MRCUT1-expressing *S. boulardii* Strain Promotes Weight Loss of PET Microplastics

#### In Vitro

To determine the capacity of our MRCUT1-expressing *S. boulardii* to degrade PET microplastics, *in vitro*, we conducted a simple weight degradation assay where pre-weighed PET discs were maintained in yeast cultures of our engineered strain adjusted to pH 6-7 with NaOH, placed on an orbital shaker over three weeks at 30°C. As a negative control, we maintained PET discs in identical media with *S. boulardii* cells expressing the SARS-CoV-2 RBD domain, which we had created for another experiment in our laboratory. The cultures were refreshed every 48–72 hours by replacing 50% of the spent medium with fresh media. Each week, the PET discs from both control and experimental groups were aseptically retrieved, re-sterilized, air-dried, and weighed to evaluate PET degradation through weight loss. All experiments were performed in triplicate.

As shown in Figure 2A, we observed a significant effect of inoculation time on the weight loss of PET plastic fragments maintained in cultures of our MRCUT1-expressing *S. boulardii* (*p* = 0.0112). Post-hoc Tukey HSD tests further indicated significant reductions in PET plastic weight between Day 0 and subsequent time points: Day 7 (*p* = 0.0122), Day 14 (*p* = 0.0020), and Day 21 (*p* = 0.0017). No significant differences were observed between later time points (Day 7 vs. Day 14, Day 7 vs. Day 21, and Day 14 vs. Day 21), suggesting that the majority of the weight loss of the PET discs occurred early during the incubation period. Linear regression analysis indicates that the weight of the PET discs decreased significantly by 17.78% ± 2.04% at day 7, 23.95% ± 7.42% at day 14, and 24.52% ± 6.81% at day 21 in the MRCUT1-expressing cells. At the same time, control samples exhibited minimal changes over the three weeks (Figure 2B). Our results suggest that we have successfully constructed a strain of the probiotic yeast, *S. boulardii*, expressing and secreting the *M. roreri* cutinase, MRCUT1, that is capable of degrading polyethylene terephthalate (PET) microplastics, *in vitro*. To the best of our knowledge, this is the first time that a cutinase capable of external PET degradation has been successfully expressed in yeast.

**Figure 2.**
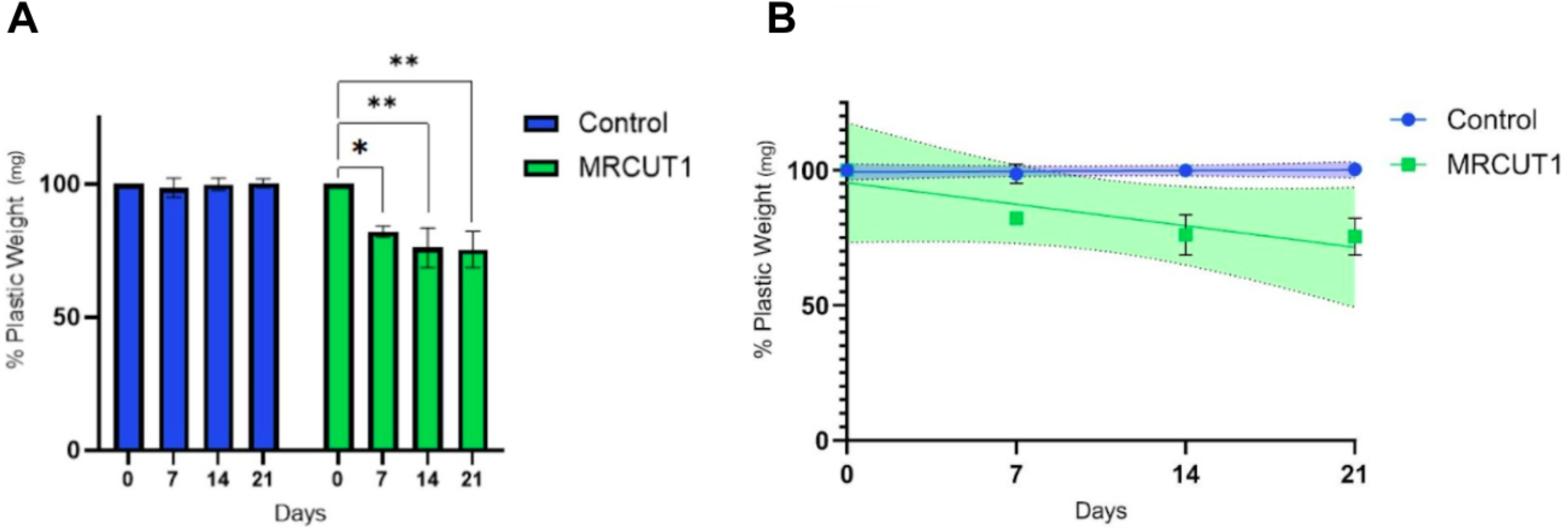
Degradation of PET Fragments by Recombinant *Saccharomyces boulardii* Cells Secreting the MRCUT1 Cutinase. (A) Bar graph comparing the percentage of remaining plastic at each time point over a 21-day period. Significant reductions in plastic weight were observed in the MRCUT1 cells (Day 0 vs. Day 7, p = 0.0022; Day 0 vs. Day 14, p = 0.0153; Day 0 vs. Day 21, p = 0.0124). (B) Linear regression of plastic weight (%) across 21 days for the Control (R^2^ = 0.1775, p = 0.6638) and MRCUT1 (R^2^ = 0.8074, p = 0.0009) groups shows significant weight loss by engineered *S. boulardii* expressing MRCUT1. Shaded areas represent 95% confidence intervals.

How does our PET-degrading *S. boulardii* strain, capable of degrading 25% of PET as measured by weight loss over three weeks, compare to its natural and engineered microbial counterparts? Numerous bacteria and fungi that naturally synthesize cutinases and PETases have been shown to degrade between 3% to 91% of PET over days, weeks, or months, depending upon the microbe tested and the technique used to measure PET loss [25]. Engineered microbes, primarily *Escherichia coli*, expressing a range of cutinases and PETases, could degrade up to 35% of PET in a month, again depending upon the microbe tested and the technique used to measure PET loss [25].

Studies like ours that measure PET degradation through polymer weight loss over 30 days, 6 weeks, 6 months, and up to 1 year have observed degradation values that did not exceed 45% [25]. As a point of comparison, when mrcut1 was expressed in *E. coli*, 31% of polyethylene terephthalate (PET) from plastic residues were degraded in a purified enzyme solution obtained from the bacteria throughout a 21-day period. No attempt was made to determine whether the bacterial cell culture could secrete the cutinase to degrade external PET polymers directly [25]. Only a few examples of recombinant bacteria that secrete engineered cutinases have been described [34,35]. In contrast, we could directly test whether PET fragments could be degraded in the presence of our recombinant yeast because we had chosen to engineer MRCUT1 so that it would be secreted into the surrounding media. This suggests that our recombinant yeast could be used for direct, external bioremediation of contaminated plastic waste. In the end, it appears that our strain’s capacity to degrade PET is comparable to that of other PET-degrading microbes, both natural and genetically engineered, with the added advantage that our strain secretes our recombinant cutinase.

### Limitations of our Study

Our study is a proof-of-concept to demonstrate that a cutinase capable of degrading PET could be expressed in and secreted by a probiotic yeast. However, our recombinant system has not been optimized. For example, we chose the commonly used TEF1 promoter to express the mrcut1 gene in *Saccharomyces boulardii* because we have had extensive first-hand experience with this promoter at the bench. However, there is evidence that suggests that other yeast promoters could better express recombinant proteins on a case-by-case basis in *S. boulardii* [36]. It is also known that wild-type cutinases, even when engineered into recombinant systems, often struggle to maintain their stability and performance over extended periods [25,37]. In principle, we could optimize the function and stability of our construct by varying the transcriptional elements in our expression system and by making targeted mutations to the amino acid sequence of our enzyme. Limited financial resources preclude us from pursuing this line of research.

Some data suggests that the pre-treatment of PET could make it more amenable to biodegradation. Physical, chemical, and biochemical methods have been adopted with varying success to alter the properties of the PET polymer, making it more accessible to microbes that secrete enzymes capable of degrading it [38]. Combinatorial strategies using PET pre-treatment (e.g., alkaline hydrolysis) might enhance substrate accessibility and accelerate enzymatic breakdown [39]. We have not attempted to explore possible pre-treatments that would improve the degradation capacity of our MRCUT1-expressing probiotic yeast strain.

### Future Research

Despite these limitations, we have created a strain of yeast that could be used for the direct, external bioremediation of environmental plastic pollution, especially in resource-limited countries.. However, we are also intrigued by the possibility that we could use our MRCUT1-expressing probiotic yeast as a prophylactic treatment to mitigate the effects of PET microplastic pollution in the intestinal tracts of common food fish, like the rabbitfish (*Siganus fuscescens*) and the milkfish (*Chanos chanos*), both of which are popular aquaculture fish in the Philippines. Hydrolytic microbes and enzymes degrade PET to bis-(2-hydroxyethyl) terephthalate (BHET) and mono-(2-hydroxyethyl) terephthalate (MHET) using PETases/cutinases. One study suggests that the chemical byproducts of PET degradation would be relatively non-toxic in fish [40]. Could feeding our MRCUT1-expressing probiotic yeast to fish in hatcheries in the Philippines decrease the detrimental effects of exposure to microplastic pollution?

## MATERIALS AND METHODS

### Yeast Strain and Culture Conditions

All experiments were conducted using the *S. boulardii* M2 strain [32], which was generously provided by Tracey J. Lamb (Emory University, USA). Cells were cultured and treated using standard yeast media and protocols, as detailed in the Cold Spring Harbor yeast handbook [41]. Unless noted otherwise, all drugs and reagents were purchased from SIGMA-Aldrich.

### Plasmid Design

The plasmid, pYEP-URA3-TEF1>α-Factor/6xHis/{mrcut1}, which was used to express and to secrete the MRCUT1 protein in yeast, was constructed and verified by VectorBuilder.com (Vector ID: B240501-112jha). The expression cassette contains the following elements: TEF1, the *Saccharomyces cerevisiae* translation elongation factor EF-1α promoter; α-Factor/6xHis/{mrcut1}, an ORF for the 6-His tagged, yeast-codon-optimized, mrcut1 gene from *Moniliophthora roreri*, with the yeast mating factor secretion signal at its N-terminus, and the CYC1 transcription terminator. A plasmid map and its sequence can be found at the VectorBuilder.com website.

### Yeast Transformation

The pYEP-URA3-TEF1>α-Factor/6xHis/{mrcut1} plasmid was transformed into the M2 *Saccharomyces boulardii* strain with the standard LiAc/SS carrier DNA/PEG method [42] with the following modification: After the initial 30 minutes of incubation at 30°C, 40μL of DMSO was added to each transformation mix. The yeast cells were then heat shocked at 42°C for 15 minutes, before being spun down and washed in sterile deionized water. Finally, the cells were resuspended in 1mL of synthetic defined (SD) media, 200μL of which was plated on SD plates lacking uracil to select for transformants. As a control, M2 *Saccharomyces boulardii* cells were transformed with a vector that drives the expression of secreted RBD from SARS-CoV-2 that was tagged with 6xHis. A map of this control plasmid can also be found at the Vectorbuilder.com website (Vector ID: VB201101-1077qmt).

### Western Blot Analysis

MRCUT1 expression in *S. boulardii* was detected via Western blotting using a standard protocol [43]. Cells were harvested after a 48-hour growth period and lysed using the Zymo Research Yeast Protein Extraction Kit. Samples were then run on a 10% Mini-PROTEAN^®^ TGX™ Precast Protein Gel (Bio-Rad) using a 10% SDS running buffer for 60 minutes at 100V. Proteins were then transferred to a PVDF membrane using a Bio-Rad Trans-Blot® SD Semi-Dry Transfer Cell at 15 V for 40 minutes. The membrane was dried at 37°C and blocked with 2% BSA for two hours at room temperature on a slow rocking platform. It was then rinsed three times for 15 minutes in 2% BSA and probed overnight with a monoclonal anti-6xHis antibody conjugated to horseradish peroxidase (Solarbio Life Sciences, K200060M-HRP) at a 1:5000 dilution in 2% BSA at 4°C. After three washes with PBST, the membrane was treated with an equal volume (1:1) mixture of ECL reagents (Solarbio, PE0011). Protein bands were detected and captured using a C-DiGit® Blot Scanner (LICORbio™).

### PET Weight Loss Assay

Approximately 1×10^6^ *S. boulardii* M2 cells transformed either with a plasmid expressing MRCUT1 or with another vector expressing the SARS-CoV-2 RBD domain as a negative control were cultured for 48 hours before the beginning of the experiment. 50μL of the saturated cultures was added to flasks containing 30mL of fresh SD-URA media adjusted to pH 6-7 with NaOH and three sterilized pre-weighed PET discs obtained from commercially available plastic bottles in the Philippines. Cultures were incubated at 30°C on an orbital shaker at 250 rpm. The cultures were replenished every 48–72 hours by replacing 50% of the spent medium with fresh media. Each week, the PET discs from both control and experimental groups were aseptically retrieved, re-sterilized, air-dried, and weighed to evaluate PET degradation through weight loss. All experiments were performed in triplicate.

### Statistical Analyses

All statistical analyses were conducted using PRISM 10. Statistical significance for the PET weight loss was determined using One-way ANOVA. Post hoc analysis using Tukey’s HSD test determined the statistical significance in the time intervals for detailed pairwise comparisons.

## ACKNOWLEDGMENTS

We thank Tracey J. Lamb (Emory University, USA) for the M2 strain of *Saccharomyces boulardii*. Our laboratory is supported by a grant-in-aid from the Philippine Council for Health Research and Development (PCHRD) of the Department of Science and Technology (DOST) of the Philippines.

## SUPPLEMENTAL MATERIAL

All data are included in this paper.

